# Revaluation of old data with new techniques reveals novel insights into the celiac microbiome

**DOI:** 10.1101/2022.10.05.510990

**Authors:** John J Colgan, Michael B Burns

## Abstract

Celiac disease is an autoimmune disorder of the small intestine in which gluten, an energy-storage protein expressed by wheat and other cereals, elicits an immune response leading to villous atrophy. Despite a strong genetic component, the disease arises sporadically throughout life, leading us to hypothesize the the microbiome might be a trigger for celiac disease. Here, we took microbiome data from 3 prior studies examining celiac disease and the microbiome and analyzed this data with newer computational tools and databases: the dada2 and PICRUSt2 pipelines and the SILVA database. Our results both confirmed findings of previous studies and generated new data regarding the celiac microbiome of India and Mexico. Our results showed that, while some aspects of prior reports are robust, older datasets must be reanalyzed with new tools to ascertain which findings remain accurate while also uncovering new findings.

**IMPORTANCE:** Bioinformatics is a rapidly developing field, with new computational tools released yearly. It is thus important to revisit results generated using older tools to determine whether they are also revealed by currently available technology. Celiac disease is an autoimmune disorder that affects up to 2% of the world’s population. While the ultimate cause of celiac disease is unknown, many researchers hypothesize that changes to the intestinal microbiome play a role in the disease’s progression. Here, we have re-analyzed 16S rRNA data from several previous celiac studies to determine whether previous results are also uncovered using new computational tools.

## INTRODUCTION

Celiac Disease (CD) is an inflammatory bowel disease (IBD) of the small intestine in which the protein gluten, found mainly in wheat and barley, causes an inflammatory response that leads to degradation of the intestinal villi (1). Left untreated, CD patients acquire a host of health problems, including malnutrition, osteoporosis and, in rare cases, cancer (1). CD is estimated to affect up to 2% of the world’s population, with Western countries having the bulk of those affected. However, many CD cases of diagnosed (1) and the disease can present at any age with a myriad of symptoms, including diarrhea, bloating, pain, nausea, insomnia, and migraines (1). Currently, the only effective treatment for CD is adhering to a gluten-free diet (GFD) (1).

The HLA DQ2 and DQ8 haplotypes are strongly associated with CD, but recent twin studies found the concordance of the disease is not 100%, suggesting environmental factors contribute to its initiation and/or progression (2). Furthermore, recent studies reported that CD subjects have dysbiotic gut microbiomes, although no clear pattern for the celiac microbiome has been defined (1). In addition, although several studies found evidence the celiac microbiome harbors an excess of potentially inflammatory bacteria, no specific bacterium or microbial community has been linked to an inflammatory response to gluten (1). The inflammatory characteristics of celiac disease could be caused or influenced by the metabolic activity of microbes, whereby byproducts of microbial metabolism modulate the immune system. This has been demonstrated for the short chain fatty-acid, butyrate, a by-product of fiber digestion by bacteria. In mice, butyrate was shown to ameliorate symptoms of rheumatoid arthritis, another autoimmune disorder, demonstrating the ability of symbiotes to modulate the immune system (3).

Other studies that examined the role of the microbiome in IBDs used the best computational tools available at the time. However, new computational tools have been developed that vastly out-perform their predecessors. For instance, previous 16S rRNA microbiome analysis was conducted using pipelines that generate operational taxonomic unit tables (OTUs). Newer computational pipelines generate amplicon sequence variants (ASVs) and produce a far more granular and accurate picture of what organisms are represented by a given sequence (4). Also, previous investigation demonstrated that ASV- and OTU-generating pipelines result in different counts of ASVs and OTUs, and that different pipelines generate distinct taxonomic assignments (5). These findings question whether conclusions drawn from older analysis methods are still relevant and whether data from previous studies needs to be reanalyzed with newer tools to obtain more insightful results.

Here, we reanalyzed data from 3 previous studies thar examined the CD microbiome. Our analysis was conducted using the dada2 pipeline to generate taxonomic classification for each read. This data was then passed off to Phangorn to create phylogenetic trees and PICRUSt2 to obtain functional analysis of the microbes. These data were then analyzed using microbiome analysis to obtain alpha- and beta-diversity metrics and identify differentially abundant taxa and metabolic pathways.

Bonder et al.(6) examined stool samples from 21 participants using the using QIIME (7), PICRUSt (8) and the greengenes database (9) to look for microbiome changes associated with the transition from a gluten-free (GF) to a gluten-containing diet (GD,. We also re-analyzed data from the study by Garcia-Mazcorro *et al*. (10), which used QIIME (7), PICRUSt (8), and Greengenes (9) to compare the stool and duodenum microbiomes of 12 celiac patients,12 non-celiac, gluten sensitive patients, and 12 controls on GD and resampled 6 months after strict adherence to the GFD. A third dataset published by Bodkhe *et al*. (11). was alsp re-analyzed. The original study included 23 untreated celiac patients, 15 first-degree relatives without celiac disease and 24 controls with hepatitis B or functional dyspepsia with paired stool and biopsy samples taken from each participant. To supplement this data, 19 healthy stool samples from Chaudhari *et al.* (12) and 17 from Dubey et al. (13) were used as controls for the analysis of stool samples from Bodkhe *et al.*. It is known that diet (14,15) and environment (16) each play a significant role in shaping the gut microbiome. From the 4 studies we were able to include 166 participants, with 31 celiac patients, 12 non-celiac gluten sensitive patients, 15 first-degree relatives, 24 patients with functional dyspepsia or hepatitis B, and 84 controls, making this work one of the largest meta-analysis examining CD across both duodenal and stool samples.

## MATERIALS AND METHODS

Studies with available data were gathered using ENA and SRA. Selected studies reported sequencing data for the v4 variable region of 16S ribosomal subunit (rRNA). Collected sequences were prepared for dada2 (7) by removing adapter sequences with cutadapt (18). Adapter sequences were reported by the parent studies. This was done for all studies except Bodkhe et al., where the adapter sequences were removed in dada2’s filterAndTrim step. Taxonomy was assigned in dada2 using the SILVA nr99 v138 training set. Phylogenetic trees were constructed using Phangorn (19, 20). The data generated by dada2 were prepared for PICRUSt2 by creating a .fasta file of the ASV sequences and a .biom table in R. See supplementary methods for a more detailed description of our analysis methods.

Data were then analyzed by PICRUSt2 (21-23) using the default parameters. We examined the metaCyc (24) pathway output of PICRUSt2. Data were then passed to microbiomeAnalyst (25, 26), where filtering was done in accordance with the methods of each respective parent study without transformation or refraction. For weighted unifrac, unweighted unifrac, Shannon diversity index, Simpson diversity index, Chao1 diversity index, RNA seq, and metagenome seq. For LEFSe, features with a P-value (unadjusted) less than 0.1 and LDA score with an absolute value of 2 or more were identified as significant.

Discrepancies were found in the control samples reported by Bodkhe et al. In response to this, additional controls were identified by searching for studies with accessible data. One alternative study was from the Delhi area of India (the same as Bodkhe et al.) and the other from Indian populations living in both rural and urban areas. The sequence data from these studies were prepared and analyzed using the protocol described above. All 3 datasets had a large number of ASVs with unassigned taxonomy. ASVs without taxonomic assignment below kingdom were tabulated using a proprietary Python script, and those with no taxonomic assignment were removed and added to a fasta file. ASVs with kingdom level assignment (bacteria) were allowed to remain. From this fasta file, 10% of the sequences were removed and clustered in mega (27). One sequence from each cluster was then BLASTed (28) to assign taxonomy to the cluster. See supplementary methods for a more detailed description of the procedures followed. Comparison in ASV assignment between Greengenes and Silva was carried out by assigning taxonomy to the Garcia-Mazcorro ASV table using both Silva nr99 v138 and Greengenes v12 databases (9, 29). The resulting taxonomy tables were then analyzed for differences using a proprietary Python script.

## RESULTS

### Reanalysis of data from Bonder et al

Bonder *et al.* had a small but significant change in alpha diversity indexes for Shannon (p = 0.0448 ANOVA) and Simpson diversity indexes (p =0.0039 ANOVA), but not for Chao1 (p = 0.2743; Supplementary Figure 1A). Samples from patients on a GFD had a higher alpha diversity compared to those on a GCD. No clustering was apparent for unifrac values (unweighted unifrac p > 0.42, weighted unifrac p < 0.0508; PERMANOVA; Supplementary Figure 1B). A single ASV was identified as being differentially abundant; this was derived from the genus *Faecalibacterium*, which was higher in patients on a GFD (Figure 1, FDR corrected p < 0.1, LEfSe LDA > 2.0).

**Figure 1.**
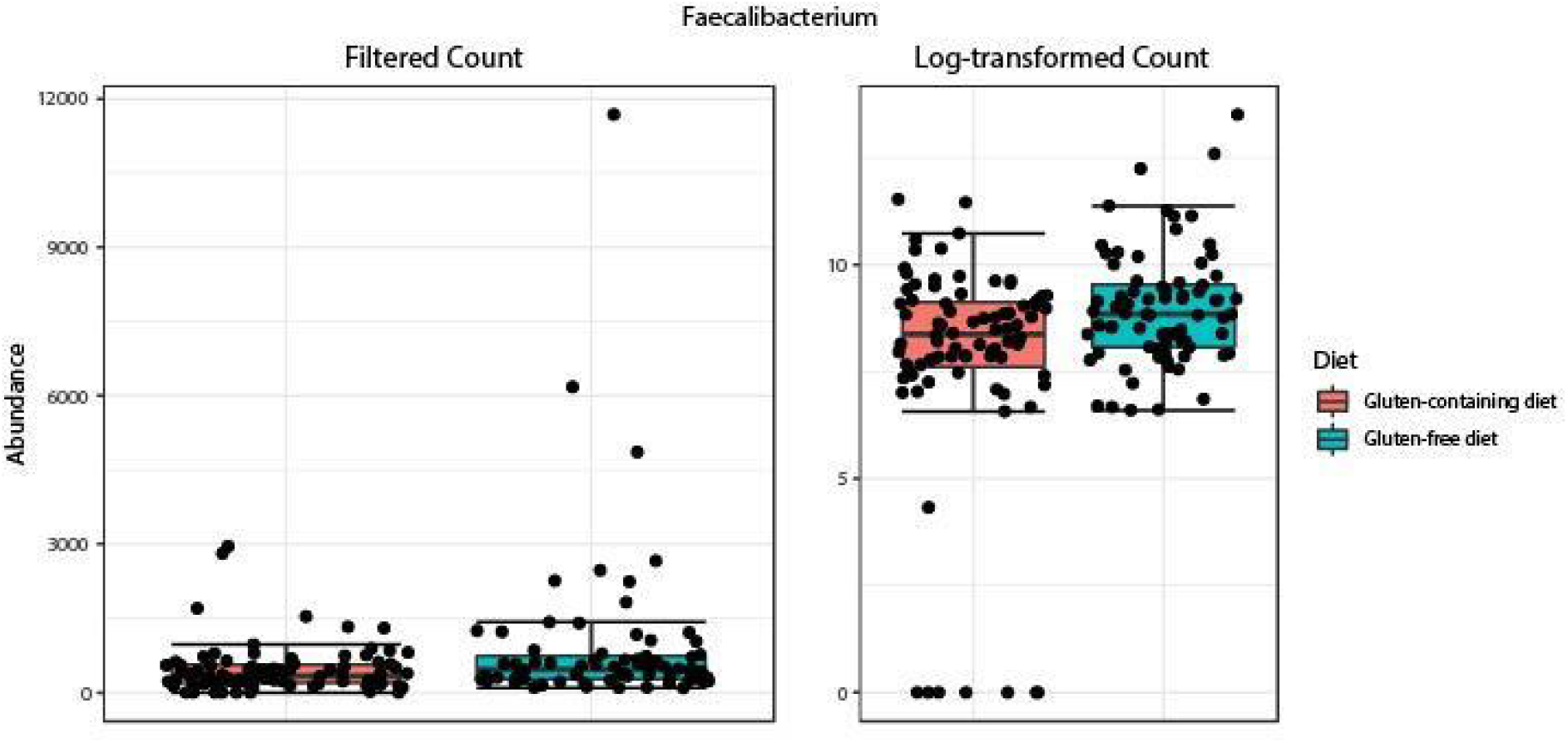
LEFSe results of Bonder *et al.*: Boxplots showing the abundance of the genus *Faecalibacterium* between GFD and GCD. The left graph shows the raw group ASV abundance and the right log-transformed abundances There was a significant increase in this ASV in the gluten-free diet study group (FDR = 0.017, LDA =2.21)

### Reanalysis of data from Garcia-Mazcorro et al

#### Duodenal analysis

Duodenum biopsies showed lowered alpha-diversity in CD patients compared to controls and NIBD, though these results were only significant for the Shannon and Simpson indixes (Supplementary Figure 2A, Chao1, p = 0.1922, Shannon, p = 0.046243, Simpson, p = 0.09176 ANOVA). No clustering was evident using unifrac (Supplementary Figure 2B, unweighted unifrac, weighted unifrac, p < 0.034, PERMANOVA). The duodenum of CD patients was characterized by elevated ASVs from *Phyllobacterium*, *Azospira*, and *Stenotrophomonas*, while the duodenum of NCGS patients had elevated ASVs corresponding to the genera *Neisseria* and *Streptococcus*. For both genuses, CD and control samples had similar average ASV counts. NCGS and control biopsies had similar levels of the genus *Fusobacterium*, but CD patients had lowered ASVs corresponding to this genus (Figure 2).

**Figure 2.**
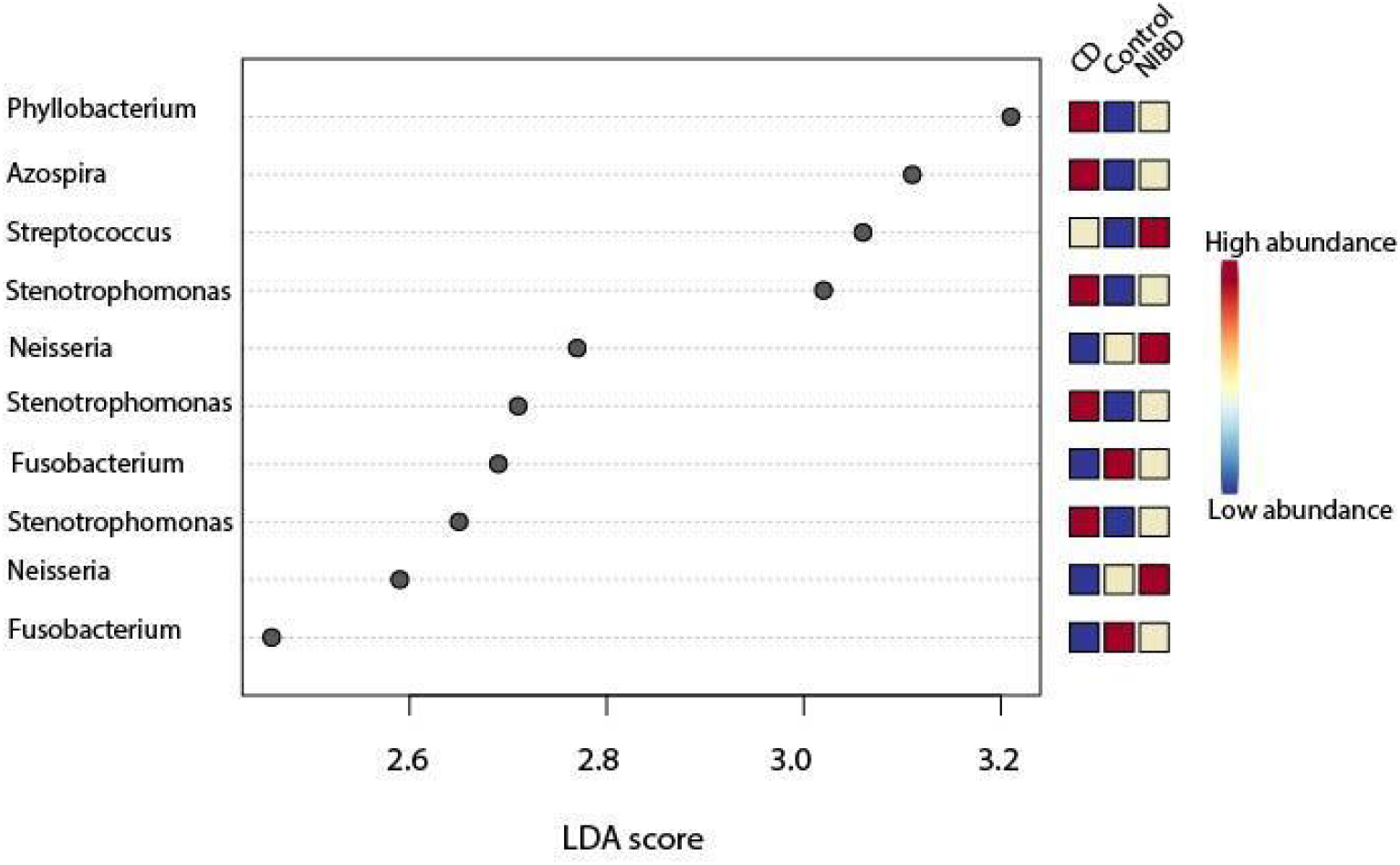
duodenal LEFSe results of Garcia-Mazcorro *et al*.: LEFSe output of differentially abundant taxa from duodenum biopsies of CD, NIBD(non-celiac gluten sensitivity) and controls. All taxa had p-values of 0.1 or less and LDA scores of 2.0 or more. It should be noted taht LDA scores are the main output of LEFSe and should be used to identify significant taxa opposed to traditional methods using adjusted or unadjusted p-values. Abundance is shown on the boxes to the right of the LDA scores with red colors indicating higher abundance relative to the other study groups and blue representing lower.

Pathway analysis showed that the microbiome of CD patients contained fewer taxa capable of menaquinone biosynthesis, with pathways for 1,4-dihydroxy-6-naphthoate biosynthesis II (PWY 7371), dimethyl menaquinone -6-biosynthesis II (PWY 7373), and menaquinone-8 biosynthesis II(PWY 6263) being lowered. All features had LEfSE LDA scores greater than 2.0 and p-values below 0.1 for RNA-seq (EdgeR 46), though not for metagenomeseq (Figure 3).

**Figure 3.**
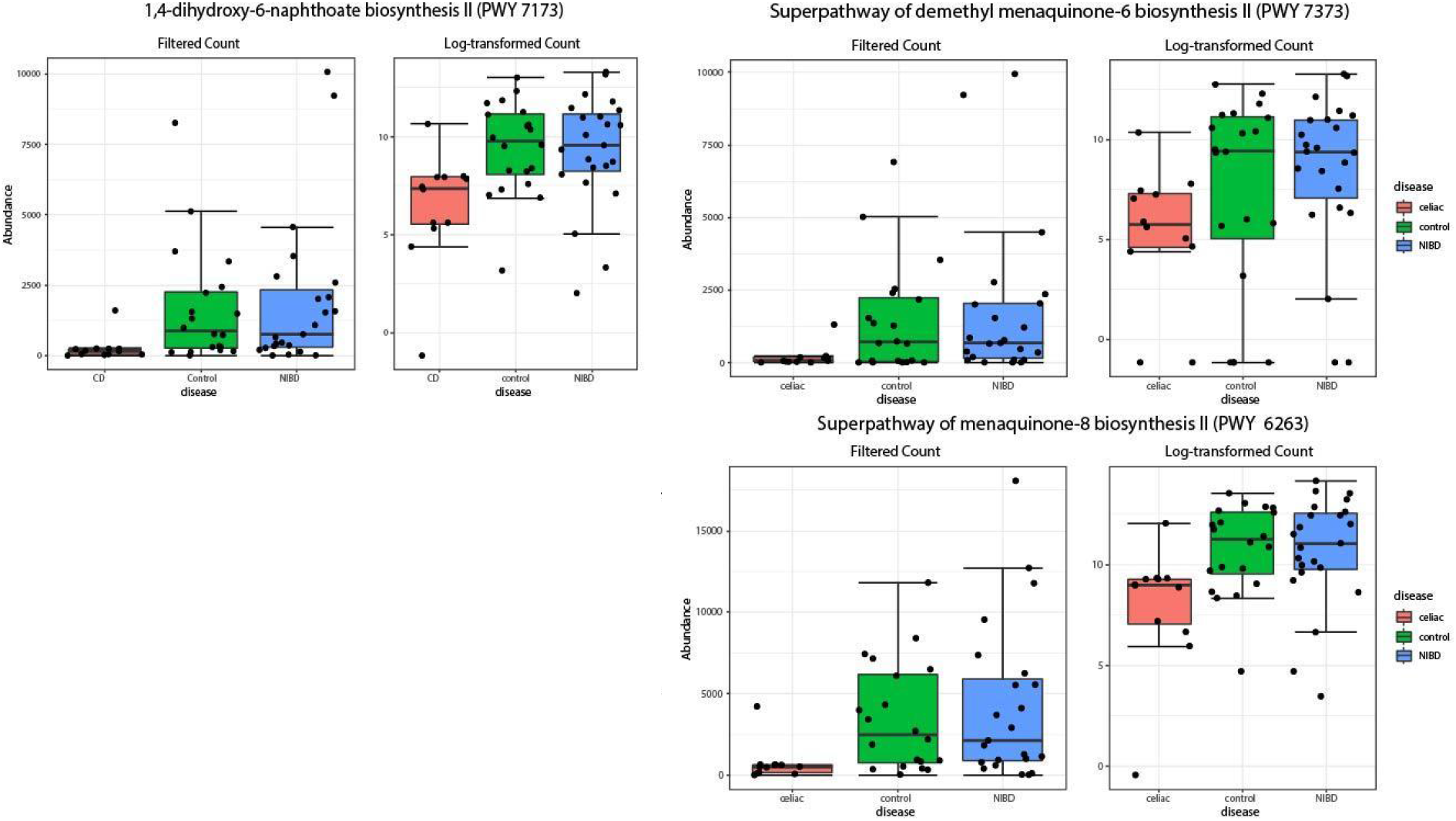
duodenal differentially abundant metabolic pathways of Garcia-Mazcorro *et al*.: Boxplots showing abundance of menaquinone biosynthesis pathways between disease states. All pathways were identified as significant using LEFSe (LDA > 2.0, p < 0.1) and EdgeR (FDR < 0.05). Each pathway was significantly reduced in CD patients compared to both healthy controls and NIBD patients. PWY 7173 had an LDA score of 3.06, a p-value of 0.0017271 and an FDR of 0.00887. PWY 7373 had an LDA score of 2.99, a p-value of 0.027261 and FDR of 0.0322283. PWY 6263 had an LDA of 3.41, a p-value of 0.0035524 and an FDR of 0.0887.

#### Fecal analysis

Fecal samples from Garcia-Mazcorro *et al.* exhibited no difference in alpha-diversity (Chao1 p = 0.79019, Shannon p = 0.61687, Simpson p = 0.58068 ANOVA, Supplementary Figure 3A). No clusters were apparent from unifrac (p < 0.145 unweighted unifrac, p< 0.179 weighted unifrac PERMANOVA, Supplementary Figure 3B). Stool samples of celiac patients had elevated levels of ASVs from *Pseudomonas* and *Novispirillum*, and lowered ASVs from *Haemophilus,* while NCGS had elevated ASVs of *Clostridia* and *Collinsella*. Control samples had an abundance of Ruminococcus and *Bifidobacterium* (p < 0.1, LDA > 2.0, Figure 4), with NCGS and CD samples having reduced ASV counts for these bacteria.

**Figure 4.**
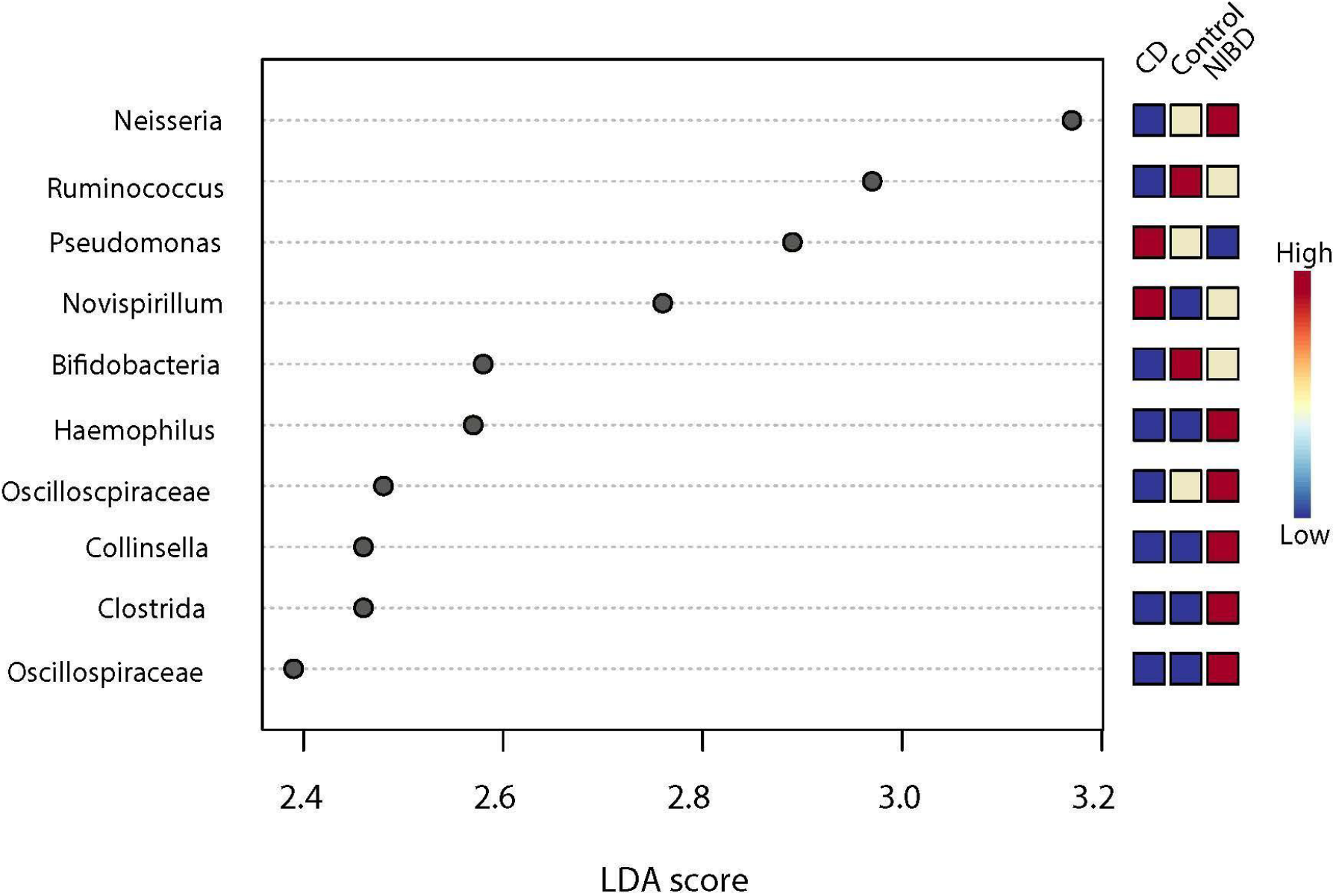
fecal LEFSe results of Garcia-Mazcorro *et al*.: LEFSe analysis of stool microbiota of Mexican healthy, NIBD and CD patients. All features had LDA scores greater than 2.0 and p-values less than 0.1. Blue colors indicate a lowered presence relative to other disease states while red colors indicate an elevated presence relative to other disease states.

114 differentially abundant pathways were identified with edgeR, but none were shared between LEFSe, metagenomeSeq, and RNA seq. However 14 pathways were shared between LEFSe and EdgeR.

#### Greengenes vs SILVA taxonomy

These data were also analyzed using the Greengenes database (version 12) to determine the extent to which pipeline choice impacted the results (9). Of the top 10 largest effect sizes producing taxa in the duodenum, 2 were assigned different taxonomy between Greengenes and Silva nr99 v138 (9, 29). ASV 14 was identified as *Clostridiales* by Greengenes but was assigned as *Stenotrophomonas* by Silva, while ASV 20 was identified as *Actinobacillus* by Greengenes v12 but as *Neisseria* by Silva (Table 1A). For stool samples, only one ASV had a mismatching assignment, with ASV 124 being identified as *Ruminococcaceae* by Greengenes and as *Oscillospriaceae* by Silva. Despite this seeming consensus among the largest effect size producing taxa, there was only a 6.3% overall similarity in ASV assignment (Table 1B). Both analyses had the same overall similarity as the ASV table used in each case was the same.

**Table 1:**
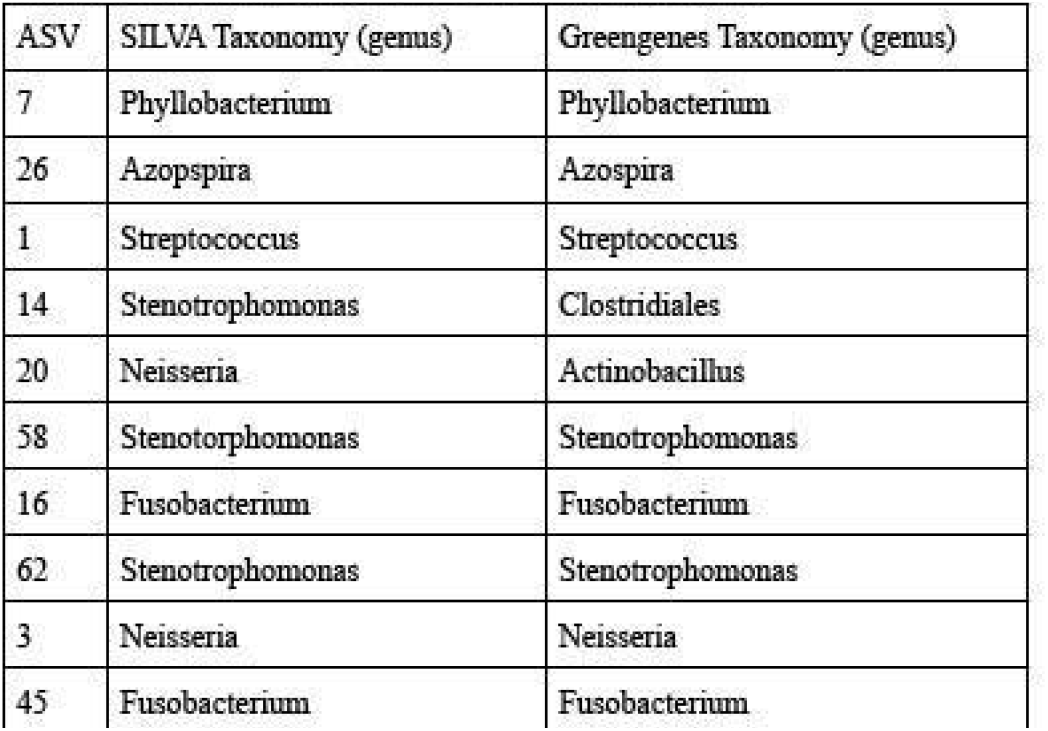
SILVA versus Greengenes Taxonomy Table 1A: Duodenum SILVA versus Greengenes Taxonomy

**Table 1B:** Fecal SILVA versus Greengenes Taxonomy

| ASV | SILVA Taxonomy (genus) | Greengenes Taxonomy (genus) |
| --- | --- | --- |
| 19 | Neisseria | Neisseria |
| 78 | Ruminococcus | Ruminococcus |
| 74 | Pseudomonas | Pseudomonas |
| 85 | Novispirillum | Novispirillum |
| 77 | Bifidobacterium | Bifidobacterium |
| 103 | Haemophilus | Haemophilus |
| 124 | Oscillospiraceae | Ruminococcaeae |
| 127 | Collinsella | Collinsella |
| 130 | Clostridia | Clostridiales (Clostridia) |
| 131 | Oscillospiraceae | Oscillospira |

### Bodkhe *et al*

#### Stool analysis

Biopsies and non-CD samples were removed, leaving just stool samples to be analyzed with controls taken from other datasets. Stool samples showed an elevated Shannon diversity index at the feature level in CD patients, however at every taxonomic level (genus through phylum) CD samples were characterized by lowered alpha diversity (Shannon index feature p = 1.0071*10^−12^ ANOVA, Shannon genus p = 3.3627*10^−31^ANOVA, Shannon family p = 6.7394*10^−28^ANOVA, Shannon order p = 3.0946*10^−29^ANOVA, Shannon Class, p = 2.9985*10^−31^ANOVA, Shannon Phylum p = 9.7104*10^−31^ANOVA, Supplementary Figure 4A). Beta-diversity analysis showed clustering for controls on the basis of region, with CD samples clustering distinctly from either control cluster (p< .05, Bray-Curtis index PERMANOVA, Supplementary Figure 4B). Unifrac also displayed clustering at the feature level for both measures (p< 0.001 PERMANOVA, Supplementary Figure 4C).

Investigation of the abundance plots revealed that the sequencing data from each study had many unassigned reads (Supplementary Figure 4D). Of the original 38,005 ASVs, 4710 had no taxonomic assignment and 11527 had only phylum level assignment. Those without taxonomic assignment were removed, leaving 33,295 ASVs. 10% of the reads without taxonomic assignment (both unclassified and without assignment below phylum) were clustered in mega, which yielded 32 clusters. 97% of the DNA sequences in the clusters corresponded to uncultured 16S bacterial DNA and 1% to contaminating human DNA. The remaining 1% was viral DNA, fungal DNA, or gDNA from *Bacteroidetes*, *Akkermansia*, and *Bifidobacterium*. Reanalysis of the data with unassigned bacteria removed (only those without taxonomic assignment) showed a similar trend as before, with the CD alpha-diversity being elevated compared to controls at the feature-level, and lowered in higher levels (Shannon feature p = 1.6529*10^−5^ ANOVA, Shannon genus p = 1.6491*10^−24^ANOVA, Shannon Family p = 9.4673*10^−21^ ANOVA, Shannon order p = 4.4937*10^−16^ANOVA, Shannon class p = 5.527*10^−10^ ANOVA, Shannon phylum p = 2.7197*10^−13^ ANOVA, p < .05; Supplementary Figure 4D and 4E), indicating the increase in diversity was due to unclassified bacteria. Weighted and unweighted unifrac also displayed results similar to those seen previously, but clustering was no longer noted at taxonomic levels genus and higher (p< 0.001, PERMANOVA, Supplementary Figure 4F).

#### LEFSe

LEFSe analysis gave mixed results, with ASVs corresponding to *Prevotella* and *Bifidobacterium* elevated or reduced levels depending upon the ASV. Healthy samples had elevated *Pseudobutyrivibrio* and *Acinetobacter*, while CD samples had an elevated ASV identified as *Bacteroidales* (Figure 5). All taxa had LDA scores larger than 2.0 and P-values below or equal to 0.1.

**Figure 5.**
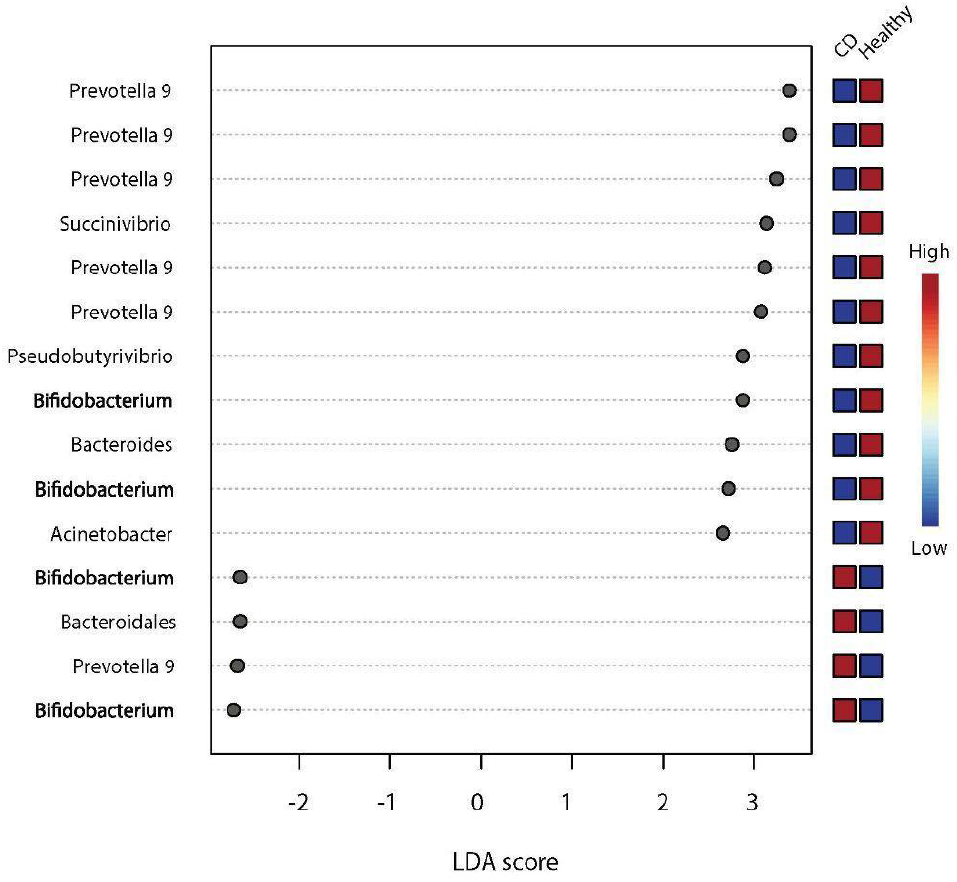
LEFSe results of Bodkhe *et al*.: LEFSe analysis of differentially abundant Indian stool staca from both healthy and CD patients. Blue indicates a lower relative abundance, while red indicates a higher relative abundance. All taxa had LDA scores with an absolute value of 2.0 or greater and p-values less than or equal to 0.1.

#### Pathways

Pathway analysis showed that samples clustered in accordance with region, with a mixed control/CD cluster forming (both sets of data taken from the Delhi region of India) and control cluster. No clustering occurred based on disease state. 69 pathways were identified as significant between edgeR, LEFSe and metagenomeSeq. The top 10 pathways with the largest effect size included anaerobic gondoate biosynthesis (PWY 7663), incomplete reductive TCA cycle (P42 PWY), L-lysine biosynthesis II (PWY 2941), cis-vaccenate biosynthesis (PWY 5973), super pathway of adenosylcobalain salvage from cobinamide II (PWY 6269) adenosylcobalamin biosynthesis from adenosylcobinamide -GDP I (PWY 5509)(Figure 10), lipid IVA biosynthesis (*E. coli*)(NAGLIPASYN), superpathway of adenosylcobalaimin salvage from coinamide I(COBALSYN PWY)(Figure 12), preQ biosynthesis (PWY 6703) and Kdo transfer of lipid IVA (*Chlamydia*)(PWY 6467) with all pathways but PWY 2941 being lowered in CD compared to controls. All pathways had an LDA score greater than or equal to 4.14 and P-values less than 0.1 (Figure 6).

**Figure 6.**
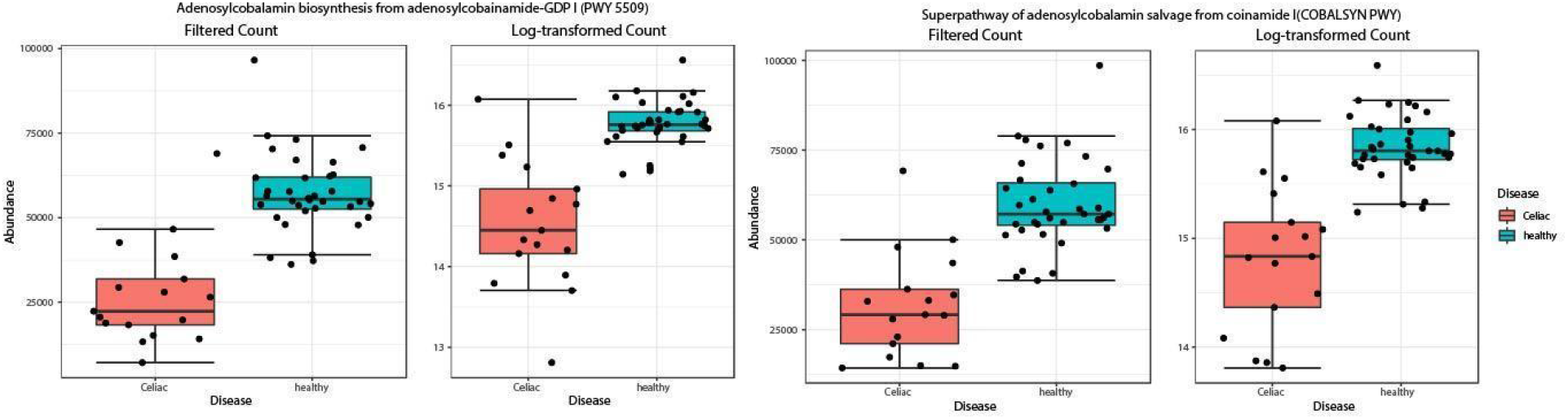
differentially abundant metabolic pathways of Bodkhe *et al*.: Boxplots showing the abundance of vitamin B producing pathways in Indian fecal samples. Plots on the left side show the abundance and log-transformed abundance of adenosylcobalamin biosynthesis from adenosylcobinamide-GDP (PWY 5509) in both CD(red) and healthy(blue) Indian stool samples. This pathway was significantly lowered in CD patients (p = 7.41817*10 ^−6^, LDA = 4.17) Similarly, the plot on the right side shows the abundance of the superpathway of adenosylcobalamin salvage from coinamide I (COBALSYN PWY) in both healthy and CD stool samples of Indian patients. This pathway was also significantly lowered in CD samples (p = 3.9879 *10 ^−6^, LDA = 4.15)

## DISCUSSION

### Impact of a GFD on healthy subjects: original analysis versus reanalysis of Bonder *et al*

Since CD patients are only offered one treatment –a GFD– it is important to separate pathologic microbiome changes that might be causative for CD from benign microbiome changes that happen due to a GFD. To control for these GFD-associated microbiome changes, we took data from a previous study that tracked microbiome changes in healthy patients eating a GFD (6). That study used QIIME, PICRUSt, and the Greengenes database for analysis and found that the transition from a GCD to a GFD resulted in negligible effects on the beta-diversity of samples, indicating the transition did not alter bacterial diversity. In contrast, our analysis detected a small but significant difference in the alpha diversity between diets, with GFD samples having a higher average alpha diversity than GCD (Chao1, Shannon, Simpson, Figure 1A).

Originally a small but significant change in beta-diversity during the transition from a GCD to a GFD was reported (Wilcoxon p-value = 0.024, weighted and unweighted unifrac, 5). PCoA analysis also showed samples tended to cluster on the basis of individual of isolation regardless of diet. Our analysis detected no differences in beta-diversity or unifrac (weighted or unweighted, Figure 1B) mirroring the original results. The original report noted shifts in the abundance of several taxa in association with diet (6), see supplementary discussion for more details. Our analysis found only one ASV that was differentially abundant, corresponding to the genus *Faecalibacterium,* LEFSe LDA >= 2.0, with a higher abundance in GFD samples (Figure 2). Both analyses found no pathways associated with the shift from a GCD to a GFD (6).

It has been noted that *Faecalibacterium*, specifically *F. prausnitzii*, are less abundant in both the untreated and treated CD microbiome (30). *F. prausnitzii* are known for producing butyrate, a short-chain fatty acid that exerts an anti-inflammatory effect by promoting the differentiation of T-cells into T-regulatory cells (31). Our work suggests the reduction in *F. prausnitzii* is unlikely due to a GFD, as it did not occur in healthy controls. Furthermore, other perturbations to the microbiome seen in treated CD patients (such as lowered alpha and beta diversity) were not noted in healthy patients on a GFD, indicating these changes are due to the disease rather than a GFD. Overall, our analysis found little impact on microbial community composition and metabolic pathways when healthy patients were placed on a GFD.

### The Mexican CD microbiome: original analysis versus reanalysis of Garcia-Mazcorro *et al*

Garcia-Mazcorro *et al*. utilized QIIME, PICRUSt, and the GreenGenes database in their analysis. Additionally, they included patients with NCGS, a condition where gluten triggers CD-like symptoms without the immune reaction and villous degradation. The original analysis noted that duodenal biopsies of CD patients had a lowered alpha-diversity (Shannon diversity index) and clustering for weighted or unweighted unifrac. We found the original results to be robust, with the only significant change being in the Shannon diversity index (ANOVA test, p = 0.046243, Figure 3A). Like the original study, no clustering was achieved with unifrac (Figure 3B).

Both studies utilized LEFSe to identify differentially abundant taxa. Originally, it was found that duodenum of CD patients was characterized by less OTUs corresponding to *Bacteroidetes* and *Fusobacteria* and more OTUs of *Novisprillium*. The microbiome of NCGS patients had elevated OTUs belonging to *Actinobacillus* and *Ruminococcaceae*, while controls had elevated OTUs for *Sphingobacterium*. Our analysis found that the biopsies of CD patients had elevated ASVs of *Azospira*, *Phyllobacterium*, and *Stenotrophomonas*. *Streptococcus* and *Neisseria* were elevated in NCGS, with both taxa having similar average group abundance in control and CD samples. Additionally, our results confirmed the deficiencies in *Fusobacteria* in CD samples found by the previous researchers (Figure 4).

Elevated *Stenotrophomonas* has been noted in other IBDs (32, 33) as well as in association with the small intestinal epithelium in mice (34). Previous studies have noted elevated *Fusobacterium* in CD (35). However, our analysis and the original analysis both showed that this genus is lowered in CD. *Fusobacterium* is considered a “bad” bacteria, as it was observed to be overly abundant in colorectal cancer, where it can inhibit T-cell-mediated immune responses and thus worsen the cancer (36). As CD is mediated by CD8+ and CD4+ T-cells (37), it is counterintuitive to expect elevated abundance of *Fusobacterium* to worsen the disease. More likely, the lowered prevalence of *Fusobacterium* upregulates T-cell mediated immune responses. Another explanation for this discrepancy is that the *Fusobacterium* detected in CD and colorectal cancer are 2 different strains, each leading to its own disease state, or that the observed changes reflect regional differences in the microbiome of Mexican CD patients, as diet and other regional factors can greatly affect microbiome composition.

*Streptococcus* and *Neisseria* were elevated in controls, with the former also elevated in NCGS and the latter in CD, highlighting a distinction between the NGCS and CD microbiomes. *Streptococcus* was previously noted as elevated in patients with NCGS (10) as well as in patients with functional dyspepsia (11). Another study noted *Neisseria* was elevated in the duodenum of Italian CD patients (38), while we demonstrated that *Neisseria* is found in similar levels in control and CD participants. This may indicate that the microbiomes of Italian and Mexican CD patients differ, or that our analysis identified a separate strain of the genus.

Previous work detected no differentially abundant pathways in the duodenum of CD, NCGS and control samples. Our re-analysis detected 3 pathways that were differentially abundant (Figure 5), PWY 7371, 7373, and 6263. All are involved in the synthesis of menaquinones (Vitamin K2), which are produced almost exclusively by gut microbes in mammals (39). This result may also explain some of the lower alpha-diversity in the CD microbiome, as menaquinones commonly serve as microbial growth factors. Furthermore, menaquinones have been shown to be growth factors of *Faecalibacterium*, perhaps explaining the previously noted deficiency for this genus in the CD microbiome (40).

Our results indicate that the CD and NCGS microbiomes are distinct, with the duodenum of CD patients being characterized by an abundance of *Stenotrophomonas* and a deficiency for *Fusobacterium,* while the duodenum of NCGS patients is characterized by an abundance of *Neisseria and Streptococcus*. Furthermore, the CD microbiome is functionally distinct as indicated by the reduced presence of 3 menaquinone-producing pathways. Reductions in these pathways may explain the known deficiencies in vitamin K among CD patients as well as the reduced alpha-diversity seen in the duodenum of these patients. Furthermore, menaquinones are known growth factors for butyrogenic bacteria and loss of menaquinone producing taxa is known to promote dysbiosis and the loss of beneficial genera of bacteria observed in CD patients (39).

Garcia-Mazcorro *et al.* obtained stool samples from patients before and after GFD treatment. However, that original study was unable to get full participation from their sample with only 5 of 12 NCGS, 6 of 12 controls and 3 of 6 CD submitting stool samples for both a GCD and a GFD. Because of this, the original authors did not focus on analyzing stool samples. Nevertheless, they did detect a shift in the proportion of *Bacteroidetes* and *Firmicutes* for all samples, regardless of disease state, with the abundance of *Bacteroidetes* being lowered across samples. No pathways were identified as being differentially abundant.

Our analysis noted no differences in alpha-diversity between study groups, as well as no differences in beta-diversity nor clustering using unifrac values (Figure 6 A/B). LEFSe identified *Pseudomonas* and *Novispirillum* as being elevated in CD stool samples, *Ruminococcus and Bifidobacterium* being elevated in controls, and *Haemophilus, Oscillospiraceae, Collinsella, Clostridia,* and *Oscillospiraceae* as being elevated in NCGS (Figure 7).

There was a difference in the identity of the significant taxa detected in the previous study and by our analysis. To understand whether this is due to pipeline (QIIME *vs.* dada2) or database (GreenGenes v12 *vs.* SILVA nr 99 v138), we assigned taxonomy to our ASV table using both GreenGenes and SILVA. We found that the most abundant taxa between Greengenes and SILVA remained 80% similar for the duodenum and 90% similar for feces (Table 1). However, the resulting taxonomy table from Greengenes and SILVA only had an overall similarity of 6.3%. This indicates that the most abundant taxa share identity between GreenGenes and SILVA, thus illustrating that the database used is most likely the cause of the discrepancies between the results of our study and the original. These results underscore the need for current researchers to reanalyze older datasets. The SRA and ENA make data freely available and easily accessible, so it is relatively simple to perform analyses like ours on older data and extract new and relevant results.

### The Indian CD microbiome: original analysis *vs.* reanalysis of Bodhke *et al*

Bodkhe *et al*. included paired biopsies and stool samples taken from 23 untreated CD patients, 24 first-degree relatives (FDRs), and 23 patients with functional dyspepsia or hepatitis B (HEPB). Their study treated FDRs as CD patients in the pre-diseased state and patients with functional dyspepsia/HEPB as controls. Our analysis aimed to compare the microbiomes of Indian CD patients to healthy controls; for that reason, FDRs and HEPB patients were left out of the analysis of the individual study. New controls were pulled from Chaudhari *et al*. and Dubey *et al*. The latter study was conducted in the same region of India (Delhi) and serves as the best control since Indian diets are regionally specific, giving different parts of the country different microbiome compositions (e.g., data from Chaudhari *et al*. was collected from a rural region of India). Together, 19 control stool samples were pulled from Chaudhari *et al.* and 17 from Dubey *et al*. No publicly available datasets containing healthy Indian duodenum biopsies were found.

The original report found no significant differences in alpha-diversity and no clustering using Bray-Curtis. Our analysis found higher alpha-diversity (Shannon index) in CD at the feature level, with alpha-diversity being lower compared to controls at all taxonomic levels (Figure 8). Beta-diversity analysis using Bray-Crustis produced 3 clusters: 2 control clusters and a diseased CD cluster. The control clusters likely reflect differences in regional diets, as both control sets were taken from different parts of India. Interestingly, CD samples clustered separately from both control sets rather than with the healthy samples from the same region, indicating differences in community structure (Figure 8B). Unweighted Unifrac showed clustering for controls and CD samples for all taxonomic levels, once again with the 3 sample groups clustering distinctly from each other.

The original report found both FDRs and CD had fewer ASVs belonging to *Dorea* and *Akermansia*. It was also noted that CD samples had a lowered abundance of *Prevotella*. FDR and CD samples had increases in ASVs corresponding to *Pediococcus, Intestinibacter, Blautia* and *Dorea* (11). Our analysis showed an elevation of *Prevotella-9* in controls with ASVs 4, 5, 6, 8, and 10. Interestingly, ASV 206 of *Prevotella-9* was elevated in CD. Healthy samples also had elevated ASVs of *Pseudobutyrivibrio, Acinetobacter,* and *Bacteroides*. *Bifidobacterium* was elevated in both CD and controls with ASV 38 being elevated in CD and ASV 50 elevated in controls. An ASV belonging to *Bacteroidales* was also elevated in CD (Figure 9).

*Prevotella* has been identified as a potentially inflammatory bacteria (41), however it has also been noted as being elevated in non-western populations, specifically in Indian populations (42) It was found that strains of *Prevotella* taken from Western and non-Western populations tend to cluster separately, with the Western populations trending toward having pro-inflammatory *Prevotella* and non-Western populations having strains of carbohydrate-degrading *Prevotella* (42, 43). The results suggest the enriched *Prevotella* in healthy Indian patients are not inflammatory. Furthermore, it was found that *Prevotella* is indeed enriched in the gut of Western IBD patients, but that these bacteria tend to be closely related to pro-inflammatory oral strains of *Prevotella* (42, 44), suggesting the ASV detected in CD patients is distinct from healthy patients. Together, these results likely show that the healthy Indian fecal microbiome is enriched in species/strains of *Prevotella* that degrade dietary carbohydrates, while the diseased Indian CD microbiome is enriched in potentially pro-inflammatory strains of *Prevotella*.

*Pseudobutyrivibrio* is a butyrate-producing bacteria that encodes many genes for plant-derived polysaccharide utilization, with butyrate being one of the end products (45). This genre was found to be lower in CD patients. From the original results of Bodkhe *et al*., *Acinetobacter* was previously noted as being lower in stool samples from Indian CD patients, Elevated abundance of *Bacteroides* was previously noted in stool of CD patients and children at risk for CD development (46). Previous studies noted a reduction of *Bifidobacterium* in the stool of CD patients (47), but we detected just one ASV of *Bifidobacterium,* perhaps reflecting species or strain differences in the *Bifidobacterium* associated with CD and controls. *Bacteroidales* were also found to be significantly reduced in patients with IBD, specifically Crohn’s disease (48). Together, these results show significant overlap between the fecal microbiomes of Indian CD patients and CD patients from around the world.

Several pathways for the salvage/production of adenosylcobalamin were identified as being reduced in the stool of Indian CD patients compared to healthy controls (COBALSYN PWY, PWY 5509, Figure 10). Adenosylcobalamin, or vitamin B12, is another critical growth factor found in microbial communities. Previous work has demonstrated that B vitamins, including vitamin B12, are widely shared in the gut microbiome, with many species lacking genes needed to produce B vitamins (49). Oral vitamin B2 supplements were shown to increase the diversity of species and ameliorate signatures of dysbiosis in fecal samples of patients with Crohn’s disease (50). Another study found deficiencies in vitamin B led to a proinflammatory state, illustrating another connection between vitamin B and its potential contributions to IBD. Furthermore, IBD symptoms were ameliorated when paired with vitamin B supplementation (51). It appears that these vitamins promote a diverse gut microbiome and the absence of B vitamin-producing bacteria and B vitamins positively correlates with worsening of IBD symptoms.

## CONCLUSION

### Reanalysis of old datasets using new tools

Older OTU-generating pipelines such as QIIME and Mothur have been used for years to conduct metagenomic studies of the gut. These tools rely on a binning approach based on a user-defined similarity threshold to denoise samples, with taxonomy assigned using a single sequence from each bin. Newer pipelines such as dada2 instead opt to use machine learning and quality scores associated with bases to denoise sequencing files and can differentiate ASVs with as little as a single base pair of difference, thus giving a far more granular picture of the microbiome.

### Metabolic differences in CD

Many of these studies also used PICRUSt1. For our analysis, we used PICRUSt2, which has a 20-fold larger database, theoretically giving it the power to derive more accurate results regarding community function. Our analysis found deficiencies in pathways for the production of electron accepting products in diseased and CD samples throughout both individual studies and pooled analysis. This trend was detected previously in dysbiotic microbiomes.

### Summary

Overall, our findings indicate that the dysbiosis observed in CD is likely a result of the disease rather than a contributing factor. Analysis of data from any geographic region individually produces results showing potentially relevant differentially abundant taxa. However no taxa were found to be differentially abundant across geographic regions.

## Supporting information

Supplemental informatio

