## Supplemental informatio for "Revaluation of old data with new techniques reveals novel insights into the celiac microbiome"

### SUPPLEMENTARY METHODS

Studies with available data were gathered using searches "celiac microbiome", "celiac disease and the microbiome", "celiac disease and gut microbiota", and "celiac disease and gut-microbiome". Studies which were selected examined the v4 variable region of the 16s ribosomal subunit (rRNA).

Collected sequences were prepared for dada2 (1). This was done using Cutadapt (2) with the command

```
cutadapt -g ADAPTERSEQUENCE1 -g ADAPTERSEQUENCE2 -o output input
```

Adapter sequences were provided by the Materials and Methods of the parent studies. This was done for all studies with the exception of Bodkhe et al., in which the adapter sequences were removed using the trimLeft = c(20,20) in dada2's filterAndTrim step. Next, the sequences were passed to dada2 (1). We used the same steps as the original paper for Bodkhe et al., since the parent study also used dada2. Both Garcia-Mazcarro et al. and Bonder et al. used single-end sequencing; adjustments were made to the pipeline in accordance with the the dada2 FAQ page for running the pipeline with single-end data. Taxonomy was assigned in dada2 using the SILVA nr99 v138 training set (3).

UPGMA phylogenetic trees for each dataset were then constructed using the R package Phangorn (4, 5) with the following commands.

```
sequences<-getSequences(seqtab.final)
alignment <- AlignSeqs(DNAStringSet(sequences), anchor=NA)
phang.align <- phyDat(as(alignment, "matrix"), type="DNA")
dm <- dist.ml(phang.align)
treeUPMA <- upgma(dm)
```

Next, the data generated by dada2 was prepared for PICRUSt2 by creating an .fasta file of the ASV sequences  and  .biom table using the following R commands with the BIOM package (6):

```
biomTable<-make_biom(t(seqtab.nochim), sample_metadata =
NULL, observation_metadata = NULL   id =
NULL,matrix_element_type = "int")
```

```
write_biom(biomTable, biom_file="table.biom")
asvTable<-seqtab.nochim
write.table(asvTable, file="ASVTableNewDataDADAbimera.txt",
row.names=TRUE , sep="\t")
write.fasta(sequences = as.list(sequences) , names =
as.list(sequences), nbchar = 80, file.out = "ASV.fasta")
```

This was then passed to PICRUSt2and run using the default parameters. The resulting data was then passed to microbiomeAnalyst (7-9) . Filtering in microbiomeAnalyst (10, 11)was done in accordance with each respective parent study's methods in mind, and no transformation or refraction was performed. For weighted unifrac, unweighted unifrac, Shannon diversity index, Simpson diversity index, Chao1 diversity index, RNA seq, and metagenome seq. For LEFSe, features with a P-value (unadjusted) less than 0.1 and LDA score with an absolute value of 2 or more were identified as significant.

Discrepancies were found with the controls in Bodkhe et al. To remedy this, new controls were found by searching for studies with accessible data using searches: "Indian microbiome" and "Indian gut-microbiota". Only studies examining the 16S v4 variable regions were used. One study from the Delhi area of India, the same as Bodkhe *et al.*, and the other from populations of Indians living in both rural and urban areas. These sequences were prepared and analyzed using the protocol described above. All three datasets had a large number of ASVs with unassigned taxonomy. To determine the identity of these sequences, ASVs without taxonomic assignment below kingdom were tabulated using a proprietary python script. ASVs with no taxonomic assignment were removed and added to a fasta file. ASVs with kingdom level assignment (bacteria) were allowed to remain. 10% of the sequences were then pulled from the resulting fasta file and clustered in mega using 95% similarity threshold (12). This threshold was chosen as it represents the variability of the v4 16S variable region. One sequence was then pulled from each cluster and assigned an identity using BLAST with default parameters(13).

Comparison in ASV assignment between Greengenes and Silva was carried out by assigning taxonomy to the Garcia-Mazcorro ASV table using both Silva nr99 v138 and

Greengenes v12 databases (14, 15). The resulting taxonomy tables were then analyzed for differences using a proprietary python script.

#### SUPPLEMENTARY DISCUSSION

#### DISCUSSION

##### GDF's impact on healthy individuals: original analysis versus reanalysis of Bonder *et al.*

**Alpha and beta diversity**

Since celiac patients are only offered one treatment–a GFD–it is important to separate pathologic microbiome changes that might be causative for CD from benign microbiome changes that happen due to the GFD. To control for these GFD-associated microbiome changes, we took data from a previous study that tracked microbiome changes in healthy patients eating a GFD (16). That study used QIIME, PICRUSt, and the Greengenes database and found that the transition from GCD to GFD had negligible impact on beta-diversity of samples, concluding that the transition did not alter bacterial diversity. In contrast, our analysis detected a small but significant difference in the alpha diversity between diets, with GFD samples having a higher average alpha diversity than GCD (Chao1, Shannon, Simpson, Supplemental Figure 1A).

**Differentially abundant taxa and functions**

Originally a small but significant change in beta-diversity during the transition from GCD to GFD was reported (Wilcoxon p-value = 0.024, weighted and unweighted unifrac, 5.). PCoA analysis also showed samples tended to cluster on the basis of individual of isolation regardless of diet, than diet. In contrast, our analysis detected no differences in beta-diversity or unifrac (weighted or unweighted, Supplemental Figure 1B) mirroring the original results. The original report noted that the species *Ruminococcus bromii* and *Roseburia faecis,* and the *Veillonellaceae* family had lowered abundance in GFD, while the families *Victivallaceae, Clostridiaceae,* and *Coriobacteriaceae*, the order *ML615J-28,* and the genus *Slackia* all increased in abundance in GFD (16). Our analysis found only one ASV that was differentially abundant, corresponding to the genus *Faecalibacterium,* LEFSe LDA >= 2.0, with a higher abundance in GFD samples

(Figure 1). The previously no significantly differentially abundant pathways were found. Likewise, we noted no differentially abundant MetaCyc pathways (17).

It has been noted that *Faecalibacterium*, specifically *F. prausnitzii*, are less abundant in both the treated and untreated celiac microbiome, than the healthy microbiome (18). *F. prausnitzii* are known for producing butyrate, a short-chain fatty acid known to exert an anti-inflammatory effect via differentiation of T-cells into T-regulatory cells (19). Our work suggests the reduction in *F. prausnitzii* is unlikely from a GDF, as it did not occur in healthy controls. Furthermore, other perturbations to the microbiome seen in treated celiac disease (such as lowered alpha and beta diversity) were not noted in healthy patients on a GFD, demonstrating that these changes are likely due to the disease rather than treatment. Overall, our findings found little impact on microbial community composition and metabolic pathways when healthy patients are placed on a GFD.

**The Mexican CD microbiome: original analysis versus reanalysis of Garcia-Mazcorro *et al.***

**Duodenal microbiome alpha and beta diversity**

Another celiac microbiome study, by Garcia-Mazcorro *et al.*, was also re-examined. That study used QIIME, PICRUSt, and the GreenGenes database and included patients with NCGS, a condition where gluten triggers symptoms similar to CD, however there is no inflammatory reaction of villous degradation unlike CD. Although duodenal biopsies of celiac patients had a lowered alpha-diversity (Shannon diversity index), they showed no differences in clustering for weighted or unweighted unifrac. Our analysis mirrored this result, with lowered alpha-diversity in celiac samples across metrics, with the only significant change being in the Shannon diversity index (Supplemental Figure 2A). Similar to the original study, there were no differences in beta diversity or clustering for unifrac values (Figure 3B).

**Differentially abundant duodenal microbiota**

The original study used LEFSe to identify differentially abundant taxa, and found the duodenum of celiac patients was characterized by less OTUs corresponding to *Bacteroidetes* and

*Fusobacteria* and more OTUs of *Novisprillium*. The microbiome of NCGS patients had elevated OTUs belonging to *Actinobacillus* and *Ruminococcaceae*. Controls had elevated OTUs of *Sphingobacterium*. Our analysis found that the biopsies of celiac patients had elevated ASVs of *Azospira*, *Phyllobacterium* and *Stenotrophomonas*. *Streptococcus* and *Neisseria* were elevated in NCGS, with both taxa having similar average group aduncance in CD and controls. *Fusobacterium* was found to be lowered in CD, with similar abundances in NCGS and controls (Figure 2).

Both *Azospira* and *Phyllobacterium* are bacteria commonly found in the roots of plants (20, 21), these findings are likely misidentifications. Elevated *Stenotrophomonas*, has been noted in other IBDs (22,23).This bacteria dominates the microenvironment near the small intestinal epithelium in dysbiotic mice (23). It is possible this bacteria is also found in close association with the gut-epithelium of the duodenum in humans, perhaps producing similar effects as those found in mice. Previous studies noted elevated *Fusobacterium* in CD (24); however, our analysis and the original analysis both showed that this genus is lowered in CD but not NCGS, illustrating a distinction between the two conditions. *Fusobacterium* is considered to be a "bad" bacteria, as it was observed as overly abundant in colorectal cancer, where, among other signaling mechanisms, it can inhibit the T-cell immune response, thus worsening the cancer (22). As CD is mediated by CD8+ and CD4+ T-cells (25), it is counterintuitive to expect elevated abundance of *Fusobacterium* to worsen the disease. More likely, the lowered prevalence of *Fusobacterium* upregulates T-cell mediated immune responses. Another explanation for this discrepancy is that it is possible that the *Fusobacterium* detected in CD and colorectal cancer are actually two different strains, each leading to its own disease state, or that the observed changes reflect regional differences in the celiac microbiome of Mexican CD patients as diet and other regional factors can greatly affect microbial composition.

*Streptococcus* and *Neisseria* were elevated in controls, with the former also elevated in NCGS and the latter in CD, highlighting a distinction between the NGCS microbiome and CD microbiome. *Streptococcus* was previously noted as elevated in patients with NCGS (26) as well as patients with functional dyspepsia (27). In the past it was noted that *Neisseria* was elevated in the duodenum of Italian CD patients (28). Here we demonstrated that *Neisseria* is found in similar levels in CD and control participants. This may indicate that the microbiome of Italian and Mexican patients differ, or previous work may have identified a different species of *Neisseria*. Together these results highlight the differences of microbiome structure between CD and NCGS.

**Differential metabolic functions: Does the celiac microbiome cause vitamin K deficiency?**

Previous work detected no differentially abundant pathways in the duodenum of CD, NCGS and controls. Our re-analysis of the pathways, detected three pathways that were differentially abundant (Figure 3). These pathways were identified as significant using both LESFe and EdgeR; and all three pathways, PWY 7371, 7373, and 6263, are involved in the synthesis of menaquinones. Menaquinones include nutrients such as vitamin K2, which are produced almost exclusively by gut microbes in mammals (29). Several case studies have noted that CD patients have vitamin K deficiencies (30, 31), however previously no explanation had been established. It is possible that these deficiencies may be due to a lowered abundance of vitamin K-producing bacteria, as indicated by the reduced abundance of pathways 7371, 7373, and 6263. This result may also explain some of the lower alpha-diversity in the celiac microbiome, as menaquinones commonly serve as microbial growth factors. Furthermore, menaquinones have been shown to be growth factors of *Faecalibacterium*, perhaps explaining the previously noted deficiency of the genus (32). A reduction in menaquinone producing taxa would result in a microbiome which is either lower in population and or diversity, either of which would result in a sample with reduced diversity.

These results indicate that the celiac and NCGS microbiomes are distinct, with the duodenum of celiac patients being characterized by an abundance of *Stenotrophomonas* and deficiency of *Fusobacterium,* while the duodenum of NCGS patients is characterized by an abundance of *Neisseria and Streptococcus*. Furthermore, the celiac microbiome is functionally distinct, as is evident by the reduced presence of 3 menaquinone-producing pathways. Deficiencies in these pathways may explain the known deficiencies in vitamin K among CD patients, as well as the reduced alpha-diversity seen in the duodenum of celiac patients. Furthermore, menaquinones are known growth factors for butyergenic bacteria; and loss of menaquinone producing taxa is known to promote dysbiosis and the loss of beneficial genera of bacteria observed in CD patients (29).

**Stool microbiota alpha / beta diversity and differentially abundant taxa**

Garcia-Mazcorro *et al.* obtained stool samples from patients before and after GFD treatment. However, that original study was unable to get full participation from their sample with only 5 of 12 NCGS, 6 of 12 controls and 3 of 6 CD submitting stool samples for both a GFD and a GCD. Because of this, the original authors did not focus on analyzing stool samples. Nevertheless, they did detect a shift in the proportion of *Bacteroidetes* and *Firmicutes* for all samples, regardless of disease state, with the abundance of *Bacteroidetes* being lowered across samples. No pathways were identified as being differentially abundant.

Our analysis noted no differences in alpha-diversity between study groups, as well as no differences in beta-diversity nor clustering using unifrac values (Supplemental Figure 3 A/B). LEFSe identified *Pseudomonas* and *Novispirillum* as being elevated in celiac stool samples, *Ruminococcus and Bifidobacterium* being elevated in controls, and *Haemophilus, Oscillospiraceae, Collinsella, Clostridia,* and *Oscillospiraceae* as being elevated in NCGS (Figure 4).

**Greengenes versus SILVA taxonomy**

There was a difference in the identity of the significant taxa detected in the previous study and our analysis. To understand whether this is due to pipeline (QIIME *vs.* dada2) or database (GreenGenes v12 *vs.* SILVA nr 99 v138), we assigned taxonomy to our ASV table, using both GreenGenes and SILVA. If our findings replicated the original study using

GreenGenes rather than SILVA, it would indicate that OTU-generating pipelines produce similar results as ASV pipelines, meaning the old analyses are valid. If the taxonomy with SILVA and GreenGenes are similar, then this would indicate the need to reanalyze older data with new tools to obtain more accurate results. We found that the most abundant taxa between greengenes and SILVA remained 80% similar for the duodenum and 90% similar for feces (Table 1). However the resulting taxonomy table from Greengenes and SILVA only had an overall similarity of 6.3%. This indicates that the most abundant taxa share identity between GreenGenes and SILVA, thus illustrating that this database is most likely the cause of the discrepancies between the results of our study and the original.

This illustrates the need of current researchers to reanalyze old datasets. Although scientists conduct analyses using best practices in their time, computational biology is rapidly evolving, with new tools and analysis techniques constantly produced. The SRA and ENA make data freely available and easily accessible, making it simple to perform analyses like ours on older data and extract new and relevant results.

#### The Indian CD microbiome: original analysis *vs.* reanalysis of Bodhke *et al.*

##### Fecal alpha/beta diversity and contaminating DNA

Bodkhe *et al*. included paired biopsies and stool samples taken from 23 untreated CD patients, 24 First-degree relatives (FDRs), and 23 patients with functional dyspepsia or hepatitis B (HEPB). Their study treated FDRs as CD patients in the pre-diseased state, and patients with functional dyspepsia/ HEPB as controls. Our analysis aimed to compare the microbiome of Indian patients to healthy controls, for that reason FDRs and HEPB patients were left out of the analysis of the individual study. New controls were pulled from Chaudhari *et al*. and Dubey *et al*. Dubey *et al*. was conducted in the same region of India (Delhi) and serves as the best control since the diets of Indians are regionally specific giving different parts of the country different microbiome compositions (e.g. data from Chaudhari *et al*. was collected from a rural region of India.) Together, 19 negative stool controls were pulled from Chaudhari *et al.* and 17 from

Dubey *et al*. No publicly available datasets containing healthy Indian duodenum biopsies were found.

The original report found no significant differences in alpha-diversity and no clustering using Bray-Curtis. Our analysis found higher alpha-diversity (Shannon index) in CD at the feature level, with alpha-diversity being lower compared to controls at all taxonomic levels (Supplemental Figure 4).

Beta-diversity using Bray-Crustis produced 3 clusters, 2 control clusters and a diseased CD cluster. The control clusters likely reflected differences due to regional diet, as both control sets were taken  from different regions of India. Interestingly, CD clustered separately from both, rather than with the healthy samples from the same region, indicating differences in community structure Supplemental Figure 4B). Unweighted Unifrac showed clustering for controls and CD for all taxonomic levels, once again with the 3 sample groups clustering distinctly from each other. Weighted unifrac showed clustering of controls and celiacs distinctly for feature level, however this pattern disappeared for genus- phylum levels (Supplemental Figure 4C). The results with Shannon diversity and weighted unifrac were puzzling as the communities should be closer in diversity for higher taxonomic levels, as opposed to the feature level. Indicating that the differences in both diversity and community structure are derived from unassigned bacterial ASVs. These ASVs are indeed bacterial in nature, as they were classified as such, indicating that better classification of non-Western microbiomes is desperately needed to better understand the contribution of the gut-microbiome in these understudied regions.

Examination of the composition of communities uncovered a high proportion of unassigned reads in the diseased state: Of the original 38,005 ASVs, 4710 had no taxonomic assignment, and 11,527 bacteria were not classified below the kingdom level, making 43% of the reads from this set uniformative (Supplemental Figure 4D). These sequences were removed, and 10% clustered by 95% similarity in mega. 95% similarity was chosen as it represents the sequence similarity of the v4 variable region and thus one cluster should encapsulate most of the potential 16S v4 sequences (33). Clustering created 32 clusters, with 1 cluster representing 96% of the sequences. One sequence from each cluster was removed and BLASTED for assignment. BLAST showed that 96% of the unclassified DNA had greatest sequence similarity to uncultured 16S rRNA records, with the remaining 3 percent split between contaminianting human, viral, fungal, and bacterial gDNA. The uncultured bacterial DNA may represent DNA chimeras; however they were not identified as chimeric using dada2's remove-chimera denovo method nor vsearch's reference-based removal method. It is likely these taxa represent organisms that have not yet been classified but are common to the Indian microbiome. Nevertheless, as these taxa exerted more of an effect on unweighted unifrac (which does not take taxon abundance into account), they are likely to only be present in small numbers within the Indian microbiome. Removing these taxa generated more robust results, with the alpha- and beta-diversity plots remaining more or less static, through the taxonomic levels, and with CD having lowered alpha diversity at the feature, genus, and family levels (Supplemental Figure 4C/D).

**Differentially abundant taxa**

The original report found both FDRs and CD had fewer ASVs belonging to *Dorea* and *Akermansia*. It was noted that CD had a lowered abundance of *Prevotella*. FDRs and CD had an increase in ASVs corresponding to *Pediococcus, Intestinibacter, Blautia* and *Dorea* (27). Our analysis showed an elevation of *Prevotella-9* in controls with ASVs 4, 5, 6, 8, and 10. Interestingly, ASV 206 of *Prevotella-9* was elevated in CD. Healthy samples also had elevated ASVs of *Pseudobutyrivibrio, Acinetobacter,* and *Bacteroides*. *Bifidobacterium* was elevated in both CD and controls with ASV 38 being elevated in CD and ASV 50 elevated in controls. An ASV belonging to *Bacteroidales* was also elevated in CD (Figure 5).

*Prevotella* has been identified as a potentially inflammatory bacteria (34), however it has also been noted as being elevated in non-western populations, specifically in Indian populations (35) It was found that strains of *Prevotella* taken from Western and non-Western populations tend to cluster separately, with the Western populations tending to have pro-inflammatory

*Prevotella* and non-Western populations having strains of carbohydrate-degrading *Prevotella* (35, 58). Non-Western populations tend to have a diet that is more rich in plant matter compared to Western populations, likely indicating that these enriched genera utilize carbohydrates from the diet (36), and that diet may explain the differences in bacterial function. Furthermore, it was found that *Prevotella* is indeed enriched in the gut of Western IBD patients, however these bacteria tend to be closely related to pro-inflammatory oral strains of *Prevotella* (35, 37). Previous studies looking at the Italian pediatric CD microbiome found that stool of CD children had deficiencies in *Prevotella* compared to non-CD children, once again suggesting that *Prevotella* may play a beneficial role in the gut (24). Together these results likely show that the healthy Indian fecal microbiome is enriched in species/strains of *Prevotella* that degrade dietary carbohydrates, while the diseased Indian CD microbiome is enriched in potentially pro-inflammatory strains of *Prevotella*.

*Pseudobutyrivibrio* is a butyrate producing bacteria that encoded for many genes for plant-derived polysaccharide utilization, with butyrate being one of the end products (38). *Acinetobacter* was previously noted as being lower in stool samples of Indian patients, via the original results of Bodkhe *et al*. Elevated abundance of *Bacteroides* was previously noted in stool of CD patients and children at risk for the development of celiac disease (24). Previous studies noted a reduction of *Bifidobacterium* in the stool of CD patients (39), but we detected just one ASV of *Bifidobacterium,* perhaps reflecting species or strain differences in the *Bifidobacterium* associated with CD and controls. *Bacteroidales* were also found to be significantly reduced in patients with IBD, specifically Crohn's disease (40).

Together these results show significant overlap between the fecal microbiomes of Indian CD patients and CD patients from around the world.

**Differential metabolic functions**

Several pathways for the salvage/production of adenosylcobalamin were identified as being reduced in the stool of Indian CD patients compared to healthy controls (COBALSYN

PWY, PWY 5509, Figure 6). Adenosylcobalamin, or vitamin B12, is another critical growth factor found in microbial communities. Previous work has demonstrated that B vitamins, including vitamin B12 are widely shared in the gut-microbiome with many species lacking genes critical for the production of B vitamins (41). Oral vitamin B2 supplements were shown to increase the diversity of species and ameliorate signatures of dysbiosis in fecal samples of patients with Crohn's disease (42). Another study found deficiencies in vitamin B led to a proinflammatory state, illustrating another connection between vitamin B and its potential contributions to IBD. Furthermore, symptoms of IBD were alemorated when paired with vitamin B supplementation (43). It appears that these vitamins promote a diverse gut microbiome and the absence of B vitamin producing bacteria and B vitamins seems to positively correlate with worsening of IBD symptoms.

A previous meta-analysis (44) of sequencing data from colorectal cancer stool and tissue samples also found that features associated with disease were not uniform across samples, but rather had a patchy distribution, with some studies having a strong signal indicating that a taxa was highly associated with the disease, while in other studies the signal was reduced or absent. To these researchers, this indicated that these taxa may be associated with the disease, and while some may worsen patient outcomes, are not a required component of the mechanism of pathogenesis in colorectal cancer. While this study was conducted on an entirely separate disease, the same may be true of CD, with some "bad" taxa associating with the disease and worsening symptoms and recovery but not actually playing a casual role in the prognosis of a patient from inactive to active CD.

39. **Olivares M, Albrecht S, De Palma G, Ferrer MD, Castillejo G, Schols HA, Sanz Y**. 2014. Human milk composition differs in healthy mothers and mothers with celiac disease - european journal of nutrition. SpringerLink. Springer Berlin Heidelberg.
40. **Gevers D, Kugathasan S, Denson LA, Vázquez-Baeza Y, Van Treuren W, Ren B, Schwager E, Knights D, Song SJ, Yassour M, Morgan XC, Kostic AD, Luo C, González A, McDonald D, Haberman Y, Walters T, Baker S, Rosh J, Stephens M, Heyman M, Markowitz J, Baldassano R, Griffiths A, Sylvester F, Mack D, Kim S, Crandall W, Hyams J, Huttenhower C, Knight R, Xavier RJ**. 2014. The treatment-naive microbiome in new-onset crohn's disease. Cell host & microbe. U.S. National Library of Medicine.
41. **Magnúsdóttir S, Ravcheev D, de Crécy-Lagard V, Thiele I**. 2015. Systematic genome assessment of B-vitamin biosynthesis suggests co-operation among gut microbes. Frontiers. Frontiers.
42. **Pham VT, Fehlbaum S, Seifert N, Richard N, Bruins MJ, Sybesma W, Rehman A, Steinert RE**. 2021. Effects of colon-targeted vitamins on the composition and metabolic activity of the human gut microbiome- a pilot study. Gut microbes. Taylor & Francis.
43. **Gominak SC**. 2016. Vitamin D deficiency changes the intestinal microbiome reducing B vitamin production in the gut. the resulting lack of pantothenic acid adversely affects the immune system, producing a "pro-inflammatory" state associated with atherosclerosis and autoimmunity. Medical Hypotheses. Churchill Livingstone.
44. **Sze M**. 2018. Leveraging existing 16S rrna gene surveys to identify reproducible ... JournalsASM. ASM Journals.

SUPPLEMENTARY FIGURES

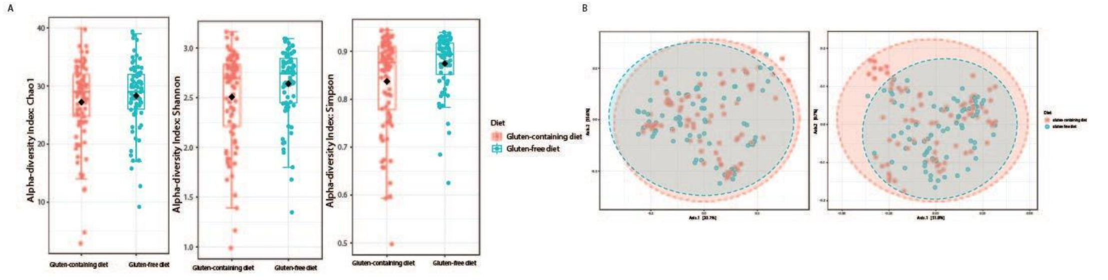


Supplementary figure 1 alpha and beta diversity analysis of Bonder et al.: A) The Chao1 alpha diversity (left) of the two study groups GFD is increased, but not significantly (p = 0.2743). Shannon diversity (middle) of the two study groups showed that GFD had a significant increase in alpha-diversity (p = 0.0448). Simpson alpha diversity index (right) showed GFD had a signifincant increase in diversity (p= 0.0039). B) Unweighted unifrac (left) and weighted unifrac (right) failed to produce clustering as a factor of diet.

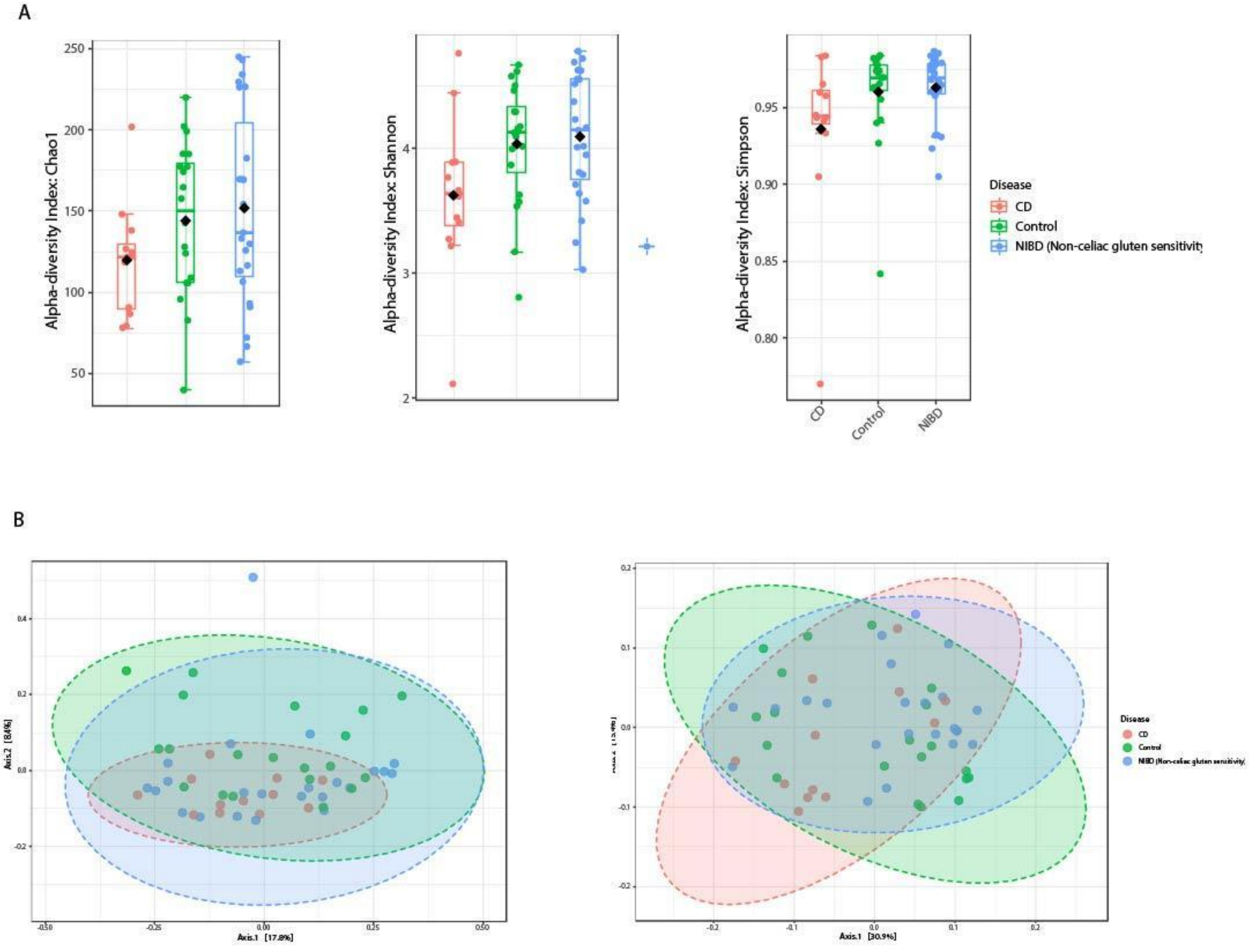


Supplementary figure 2 duodenal alpha and beta diversity analysis of Garcia-Mazcarorro et al. A) Alpha-diversity analysis of duodenum samples. No test was significant (P> .05). B) Unifrac analysis depicting clustering ofsamples based on community structureand relatedness of samples. Neither weighted unifrac (right) nor unweighted unifrac (left) were able to produce accurate clusters for CD, NIBD, and controls.

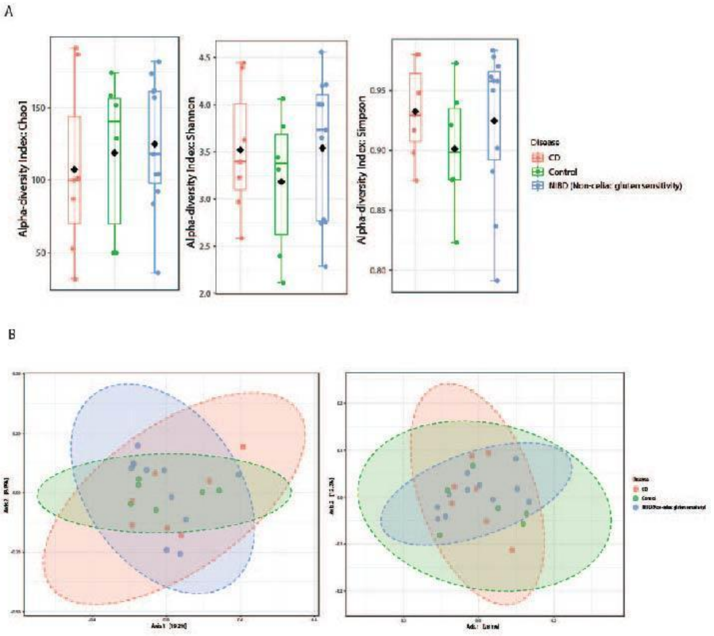


Supplementary figure 3 fecal alpha and beta-diversity analysis of Garcia-Mazcorro et al.: A) Alpha-diversity analysis of stool samples. No significant difference was noted in Chao1, Shannon or Simpson diversity indexes (p > 0.05). B) Weighted unifrac (right) and unweighted unifrac (left) were unable to produce clusters based on disease.

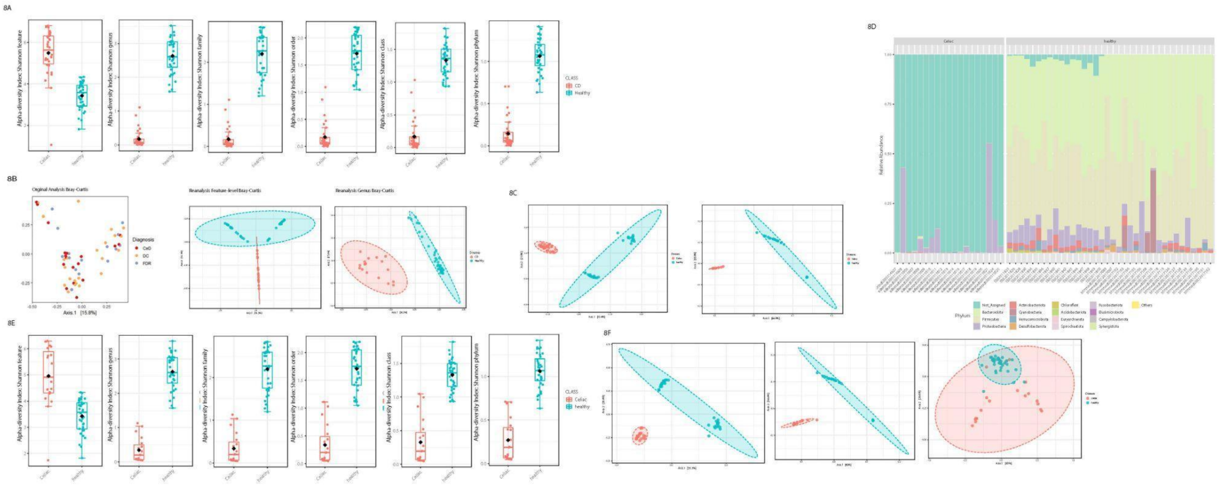


Supplementary figure 4 alpha and beta diversity of Bodkhe et al A) Shannon diversity index the from genus to phylum level. CD patients had enriched stool diversity at the feature level, while controls had increased diversity at the genus through phylum levels (Shannon index feature P = 1.0071 *10-12, Shannon genus P = 3.3627*10-31, Shannon family P = 6.7394 *10-28, Shannon order P = 3.0946*10-29, Shannon Class, P = 2.9985*10-31, Shannon Phylum P = 9.7104*10-31). B) Original analysis of Bray-Curtis index versus reanalysis at the feature and genus level. Reanalysis showed distinct clustering at feature and genus levels (P< 0.01). C) Unifrac unweighted (left) and weighted (right) analysis of CD and non-CD stool samples showed distinct clustering (P<0.001). D) Stacked area abundance bar plots at the phylum level. CD samples were characterized by a large degree of unassigned taxa. E) Shannon diversity of stool samples with unassigned taxa (Kingdom level removed). CD samples had increased alpha diversity at the feature level but loweralpha diversity at all other taxonomic levels (Shannon feature P = 1.6529*10-5, Shannon genus P = 1.6491*10-24, Shannon Family P = 9.4673*10-21, Shannon order P = 4.4937*10-16, Shannon class P = 5.527*10-10, Shannon phylum P = 2.7197*10-13). F) Unweighted unifrac (right) and weighted unifrac (feature-level middle, genus right) analysis. Weighted unifrac only produced clusters at the feature level while unweighted did for all taxonomic levels (P< 0.001).

### SUPPLEMENTARY FIGURE LEGENDS

Supplementary figure 1 alpha and beta-diversity analysis of Bonder *et al.*: A) Chao1 alpha-diversity (left) of the two study groups GFD is increased but not significantly (p = 0.2743). Shannon diversity (middle) of the two study groups showed that GFD had a significant increase in alpha-diversity (p - 0.0448). Simpson alpha-diversity index (right) showed that GFD had a significant increase in diversity (p = 0.00393). B) Unweighted unifrac (left) and weighted unifrac (right) failed to produce clustering as a factor of diet.

Supplementary figure 2 duodenal alpha and beta-diversity analysis of Garcia-Mazcorro *et al.*: A) Alpha-diversity analysis of duodenum samples. No test was significant (p > 0.05) B) Unifrac analysis depicting clustering of samples based on community structure and relatedness. Neither weighted unifrac (right) nor unweighted unifrac (left) were able to produce accurate clusters for CD, NIBD and controls.

Supplementary figure 3 fecal alpha and beta-diversity analysis of Garcia-Mazcorro *et al.*: A) Alpha-diversity analysis of stool samples. No significant difference was noted in Chao1, Shannon or Simpson diversity indexes (p > 0.05). B) Weighted unifrac (right) and unweighted unifrac (left) were unable to produce clusters based on disease.

Supplementary figure 4 alpha and beta-diversity of Bodkhe *et al.*: A) Shannon diversity index from the genus to phylum level. CD patients had enriched stool diversity at the feature level, while controls had increased diversity at the genus through phylum levels (Shannon index feature p = 1.0071 *10 -12, Shannon genus p = 3.3627*10-31, Shannon family p = 6.7394*10-28, Shannon order p = 3.0946*1--29, Shannon class = - 2.9985*10-31, Shannon phylum p = 9.7104*10-31). B) Original analysis of Bray-Curtis index versus reanalysis at the feature and genus level. Reanalysis showed distinct clustering at the genus and feature levels ( p < 0.01). C) Unifrac unweighted (left) and weighted (right) analysis of CD and non-CD stool samples showed distinct clustering (p < .001). D) Stacked area abundance plots at the phylum level. CD samples were characterized by a large degree of unassigned taxa. E) Shannon diversity of stool samples with unassigned taxa (kingdom level) removed. CD samples had an increased alpha-diversity at the feature level but lowered alpha-diversity at all other taxonomic levels (Shannon feature p = 1.6529*10-5, Shannon genus p = 1.6491*10-24, Shannon family p = 9.4673*10-21, Shannon order p = 4.49317*10-16, Shannon class p - 5.527*10-10, Shannon phylum p = 2.7197*10-33) F) Unweighted unifrac (right) and weighted unifrac (feature-level middle, genus, right) analysis. Weighted unifrac only produced clusters at the feature level while unweighted did for all taxonomic levels (p < 0.001).
